## Supplemental Material for "A Collaborative Multi-Model Ensemble for Real-Time Influenza Season Forecasting in the U.S"

### Supplement for
### "A Collaborative Ensemble Approach to Real-Time Influenza Forecasting in the U.S.: Results from the 2017/2018 Season"

Nicholas G Reich, Craig McGowan, Teresa Yamana, Abhinav Tushar, Evan L Ray,
Dave Osthus, Sasikiran Kandula, Logan Brooks
Willow Crawford-Crudell, Graham Casey Gibson, Evan Moore, Rebecca Silva
Matthew Biggerstaff, Michael A Johansson, Roni Rosenfeld, Jeffrey Shaman

February 11, 2019

#### 1 Forecast accuracy by region and target

See Figure 1.

#### 2 Supplemental evaluation metrics

##### 2.1 Bias and MSE

We compared the `FSNetwork-TTW` ensemble model's accuracy in 2017/2018 to the top-performing models from each team in the training phase. We used the metrics of root mean-squared error (RMSE) and average bias to measure accuracy of point estimates. Note that the ensemble weights were optimized solely to maximize log-score, so these accuracy scores are not an indicator of how well the ensemble could do if it were optimized to minimize point-estimate error. Consistent with the CDC scoring rules, we only evaluated point estimates within the "scoring bounds" specific to each target, region, and season (see Methods in main manuscript).

Overall, the `FSNetwork-TTW` model ranked second in both RMSE and average bias, behind the `LANL-DBM` model (Figure 2). All selected models showed a negative bias (i.e. underestimation, on average) of the targets on the wILI scale (week-ahead incidence and peak percentage). The `CU-EKF_SIRS` model showed particularly low bias for 1- and 2-week-ahead foreacasts, although greater variability led to lower ranks for RMSE.

In general, these results suggest that using separate weighting schemes for point estimates and predictive distributions may be valuable.

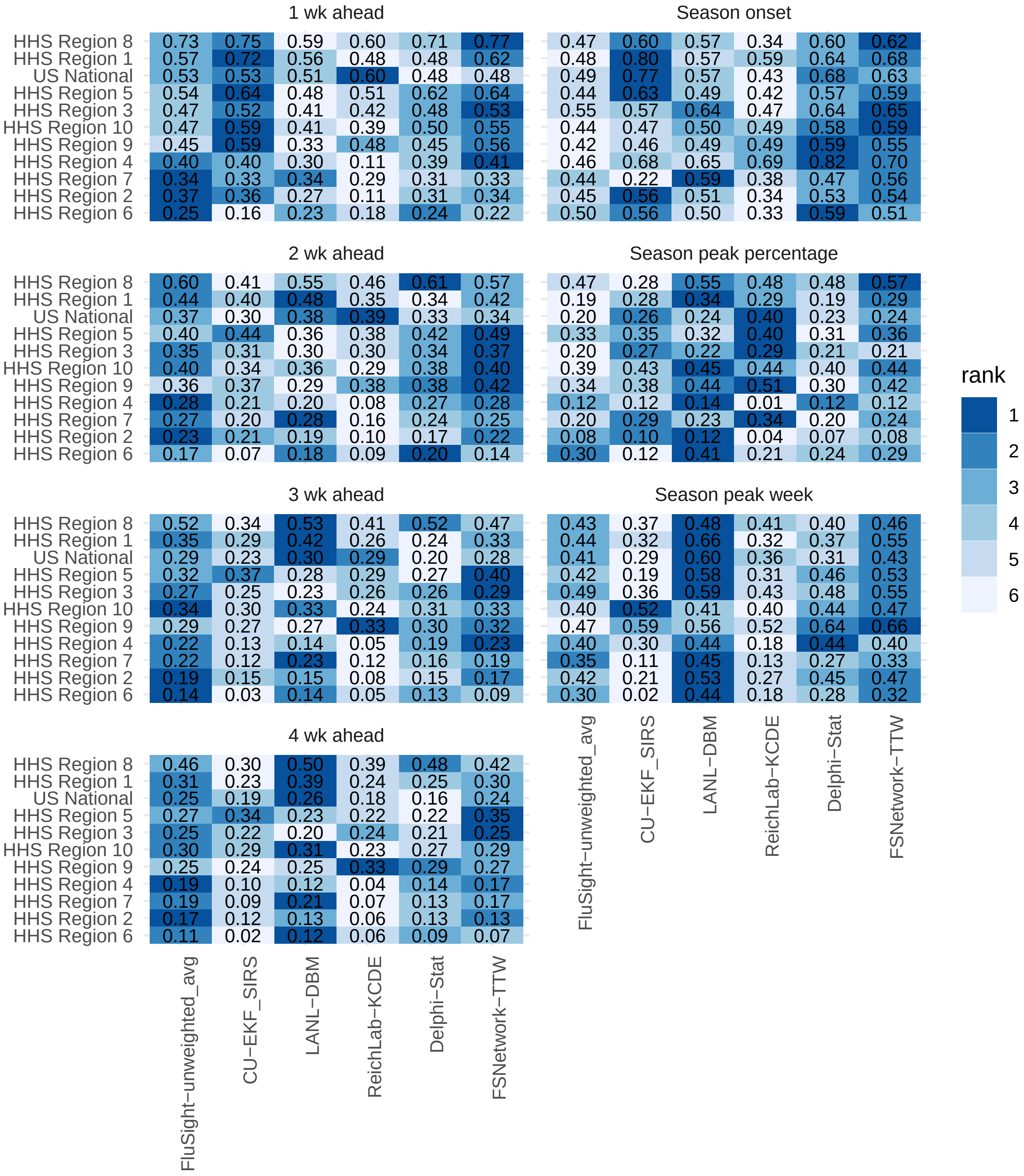


Figure 1: Average forecast scores and ranks by target and region for 2017/2018. Only selected models are shown. Regions are sorted with the most predictable region overall (i.e. highest forecast scores) at the top. Color indicates model rank in the 2017/2018 season (darker color indicates better average score). The average forecast score is printed.

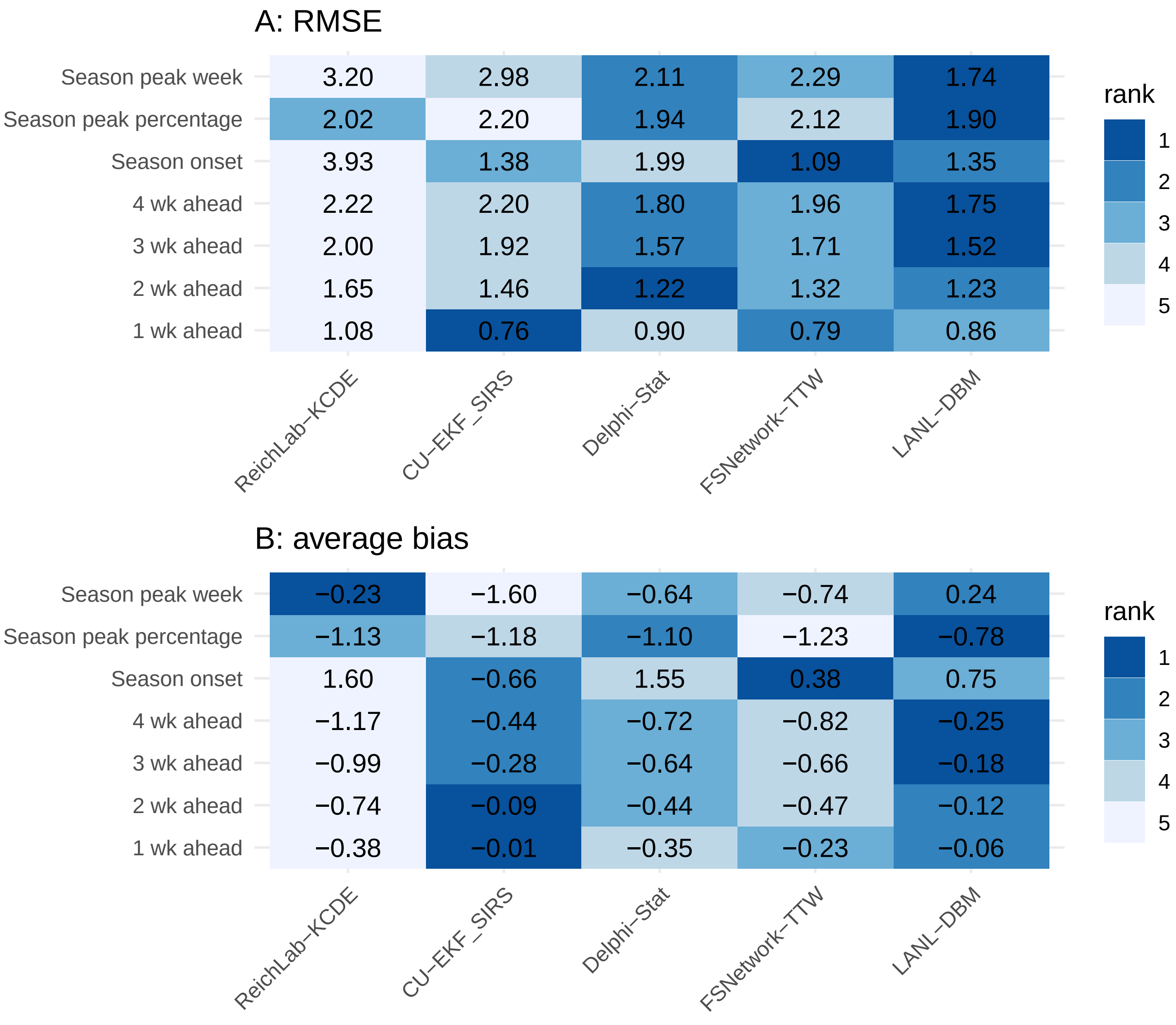


Figure 2: Root mean squared error (RMSE, panel A) and bias (panel B) by target for selected models (with rank) in the 2017/2018 season. Evaluations are for all weeks in the 2017/2018 season. Models are sorted with lowest RMSE on right.

#### 2.2 Probability integral transform

The Probability Integral Transform (PIT) is an evaluation metric that can be used to assess the calibration of a predictive model. A common application of PIT is testing whether a set of values from an unknown target distribution can be accurately modeled by one or more predictive distributions. In brief, statistical theory tells us that if we plug the set of observed values (which come from the unknown target distribution) into the cumulative distribution function of the predicted distribution, the output, otherwise known as the PIT values, should be uniformly distributed if the predicted distribution matches the true distribution.[1, 2] Therefore, looking at the PIT values provides a quantitative and qualitative assessment of the predictive model calibration by comparing the shape of the histogram of PIT values to a uniform distribution. Intuitively, the PIT measures how often a model's probabilistic assessment is true, i.e. does an event that the model says has a 10% chance of occuring really only occur 10% of the time. Systematic deviations from the expected uniform distribution may indicate lack of calibration in some aspects of the predictive model.

In our application, we evaluate the set of all probabilistic forecasts of the five targets on the wILI scale (1 through 4 week-ahead wILI percent and the peak percentage) from the `FSNetwork-TTW` model using PIT. For the 2017/2018 influenza season, we obtained a PIT value from each predictive distribution based on region, target, and week of season. As in other evaluations presented here, we only considered forecasts from the time-period of interest for the CDC, depending on the timing of the peak for each region-season. We rounded each PIT value to the nearest tenth of a decimal place and plotted them on a historgram with ten bars, one for each decile of the Uniform(0,1) distribution. We computed a Monte Carlo confidence interval under the null hypothesis that the PIT values are independent and identical draws from a Uniform(0,1) distribution, conditional on the number of PIT values.

In the 2017/2018 season, the `FSNetwork-TTW` model showed good calibration for all week-ahead targets (Figure 4). The models appeared to be slightly better calibrated for shorter forecast horizons (i.e. 1- and 2-week ahead) than for longer horizons. Forecasts for the peak percentage were less well calibrated, with more forecasts than expected occuring in both low and high tails of the predictive distribution.

Across all training seasons, the `FSNetwork-TTW` model showed some lack of calibration for all targets considered (Figure 3). In particular, the predictive distributions appeared to be too wide, with eventually observed values falling in the central region of the distribution more than expected. A slight negative bias is evident as well, from the skewness of the PIT figures, with the observed values more likely to fall under the median of the distribution for 2-, 3-, and 4-week-ahead forecasts. However, these forecasts were evaluated on only seven seasons worth of data, and given that the forecasts were better calibrated in a larger-than-usual season such as 2017/2018 (Figure 4) suggests that the model in the training phase may have been appropriately allowing for the possibility of such a large season.

All in all, there could be some improvement in model calibration as measured by PIT. However, to date, this has not been designated as an explicit target for optimization of the ensemble weighting schemes.

### 3 Updated weights incorporating 2017/2018 performance

See Figure 5.

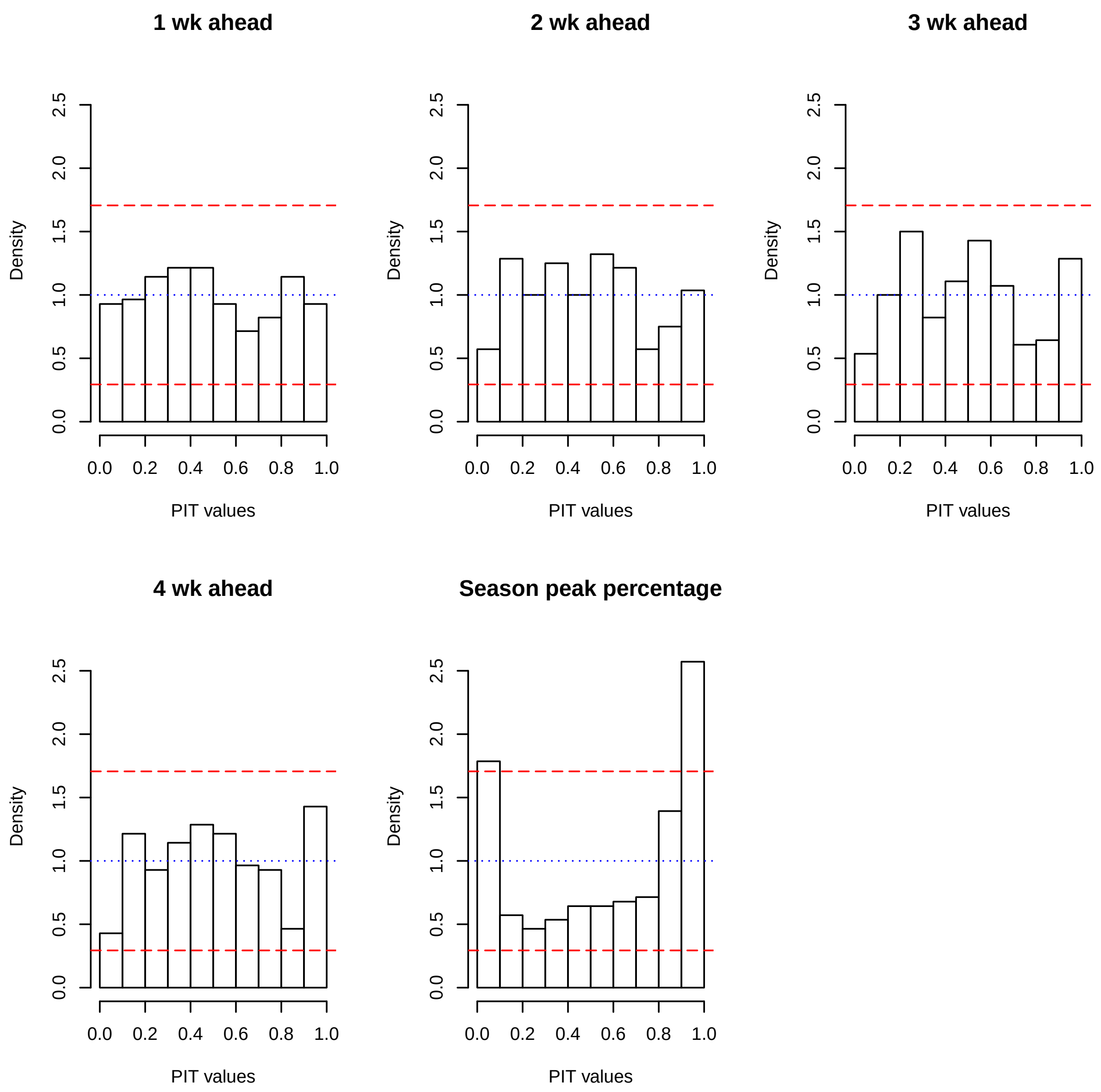


Figure 3: Probability integral transform histograms by target for the FSNetwork-TTW model in the 2017/2018 season. If the model is well-calibrated, the histogram should resemble a uniform distribution, i.e. all the bars should be level at $y$=1. The red dashed lines represent a Monte Carlo confidence interval under the null hypothesis that the PIT values follow a Uniform(0,1) distribution. The fact that some of the bars for the season peak percentage target lie outside the CI bounds suggest that the model shows significantly weak calibration for that particular target.

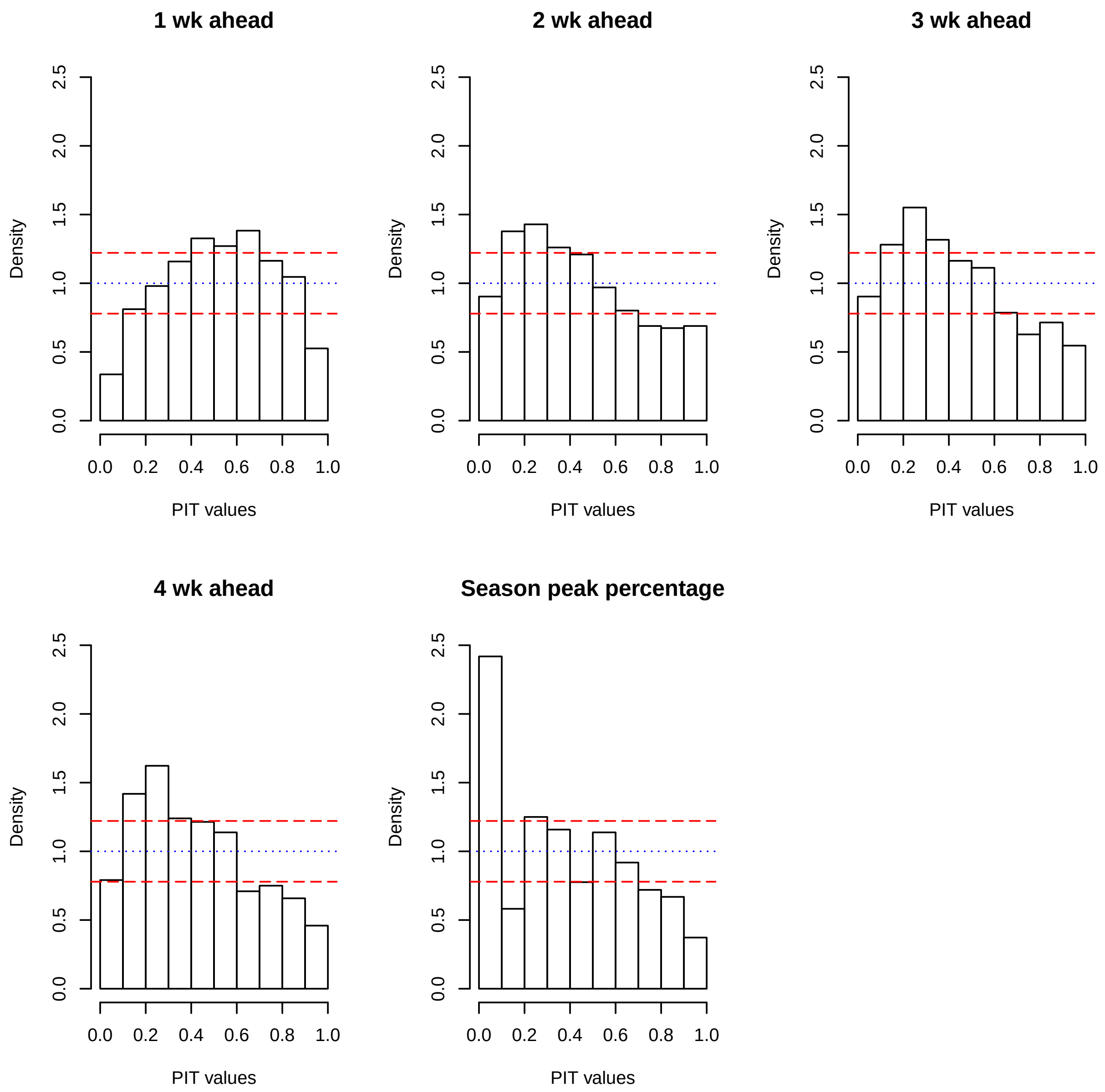


Figure 4: Probability integral transform histograms by target for the FSNetwork-TTW model in all training seasons: 2010/2011 through 2016/2017. If the model is well-calibrated, the histogram should resemble a uniform distribution, i.e. all the bars should be level at $y$=1. The red dashed lines represent a Monte Carlo confidence interval under the null hypothesis that the PIT values follow a Uniform(0,1) distribution. The intervals are narrower here than in Figure 3 because there are more observations from all the training seasons combined than in the one testing season. The model is showing some significant lack of calibration for all targets. In particular for the season peak percentage, substantially more observations were in the lowest 10% of the predictive distributions than would have been expected due to chance.

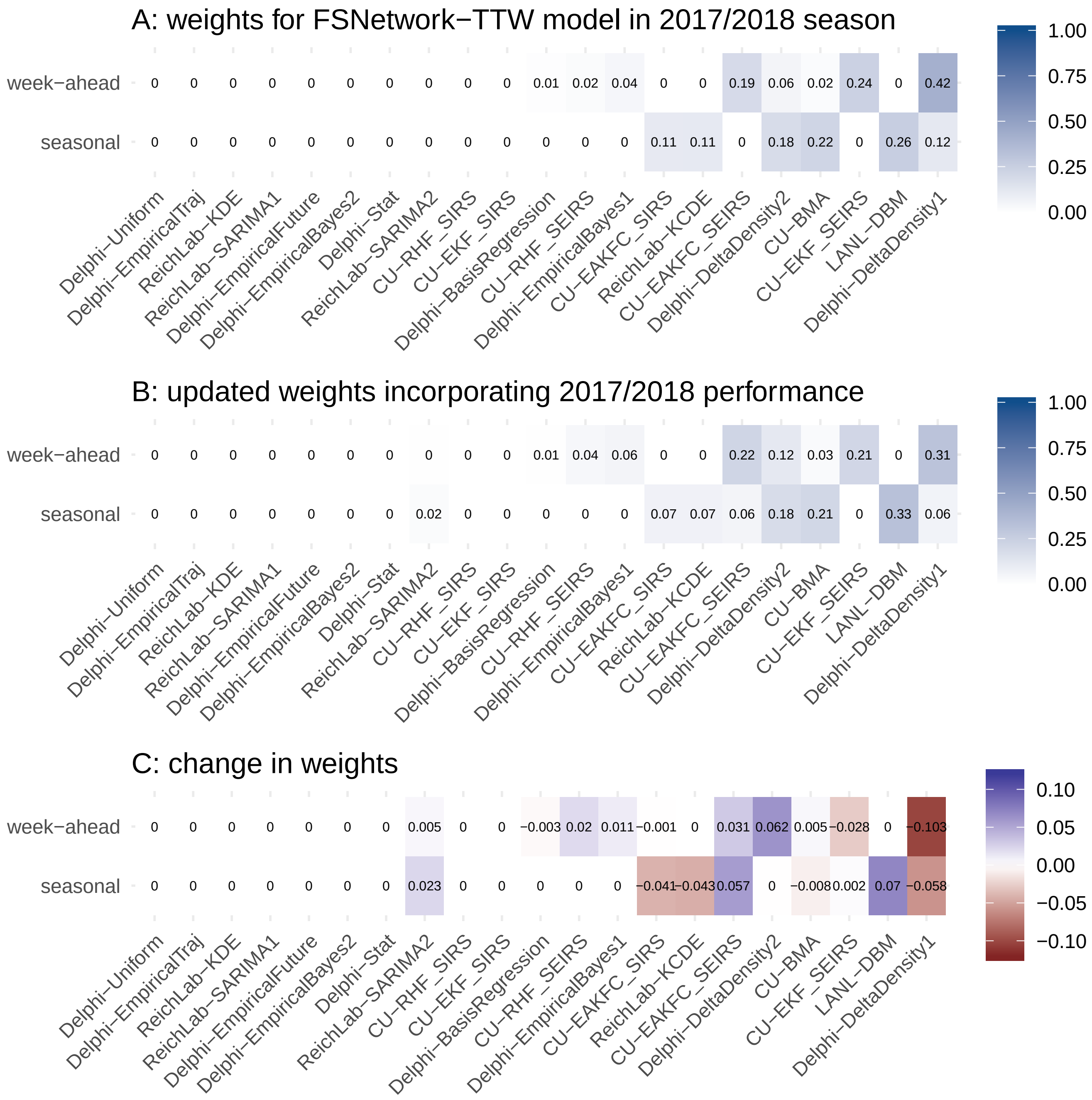


Figure 5: The change in the estimated weight for each model after including the 2017/2018 performance results.

### 4 EM Algorithm for Weighted Density Ensembles

The following is adapted from [3, 4] and describes the use of the Expectation Maximization (EM) algorithm for constructing weighted average ensemble models in the context of infectious disease forecasting. Our goal is to develop a weighted density ensemble that combines the full predictive distributions in such a way as to optimize the score of the resulting model average.

To use the EM algorithm to find optimal weights, we formulate the question as a missing data problem. We consider a data generating process in which an observed target is generated from $f(z)$ by choosing one of the $f_c(z)$ component distributions as a random draw from a multinomial distribution with probabilities $\pi_c$. Here we supress the subscripts for target, region and week ($t$, $r$, and $w$) for simplicity. The problem is that we do not know, for each observed datapoint $z_i^*$, which component this observation was drawn from. However, we can make a best guess, conditional on the data and our current estimates of $\pi_c$, of how often each component was chosen. This is the "E step". Then, based on these guesses, we can update our estimate of $\pi_c$. This is the "M-step".

The "E step" of the EM algorithm we can think of as determining, for each component $c$, the expected number of times for each of our observed $N$ datapoints that component $c$ was chosen as the contributor to $f(z)$:

$$\mathbf{E}[model_c|data] = \sum_i \frac{\pi_c f_c(z_i^*)}{f(z_i^*)} \tag{1}$$

Heuristically, we can think of the expression $f_c(z)$ equivalently as $Pr(z|modelc)$ or in words the likelihood of seeing the value $z$ given that component $c$ is the "chosen" model.

The "M step" of the EM algorithm simply calculates, conditional on the "complete data", i.e. the $z_i^*$ and the estimated number of times each component was chosen, the fraction of times each method was chosen. Therefore, if we simply divide the quantity from the "E-step" by $N$, our total number of observations, we obtain a new estimate of this probability:

$$\pi_c^{(k+1)} = \frac{1}{N}\mathbf{E}[model_c|data] \tag{2}$$

$$= \frac{1}{N}\sum_i \frac{\pi_c^{(k)} f_c(z_i^*)}{f(z_i^*)} \tag{3}$$

Assume that we have a set of $C$ fitted predictive densities "evaluated at" observed data $z_i^*$ for $i = 1, ..., N$. In our application, we let the $f_c(z_i^*)$ be computed as the probabilities associated with the modified scores as described in the main manuscript. As an example, for season peak percentage and the short-term forecasts, probabilities assigned to wILI values within 0.5 units of the observed values are included as correct, so the modified score becomes $f_c(z_i^*) = \int_{z_i^*-.5}^{z_i^*+.5} f_c(z|\mathbf{x})dz$. We will notate these scores as $f_c(z_i^*|\mathbf{x})$. There will be $C \cdot N$ total observations, as each model must have an associated score (a probability, between 0 and 1) for each observed data point.

We wish to obtain a set of optimal weights $\tilde{\pi} = \{\tilde{\pi}_1, \tilde{\pi}_2, ..., \tilde{\pi}_C\}$ for combining the models such that $\forall c\ \tilde{\pi}_c \geq 0$ and $\sum_{c=1}^{C} \tilde{\pi}_c = 1$. The weights can be used to then combine the component models into an ensemble model as

$$f(z|\pi) = \sum_{c=1}^{C} \pi_c f_c(z).$$

We define a function $\ell(\pi)$ that computes a log-likelihood of the resulting ensemble as follows:

$$\ell(\pi) = \frac{1}{N} \sum_{i=1}^{N} \log f(z_i|\pi).$$

Below, we define one procedure to obtain a set of weights for the ensemble.

**Algorithm 1** Degenerate Expectation Maximization (DEM) algorithm

1: **procedure** DEM(...)
2: Initialize $\pi_c^{(0)}$ such that $\forall c$ $\pi_c^{(0)} \geq 0$ and $\sum_{c=1}^{C} \pi_c^{(0)} = 1$
3: Set $t = 0$
4: Set $\Delta = 1$, or another arbitrary constant.
5: Set $\epsilon$ to be a very small positive number strictly less than $\Delta$.
6: **while** $\Delta > \epsilon$ **do**
7: Set $t = t + 1$
8: Update weights, $\forall c$, $\pi_c^{(t)} = \frac{1}{N} \sum_{i=1}^{N} \frac{\pi_c^{(t-1)} f_c(z_i)}{f(z_i|\pi^{(t-1)})}$
9: Set $\Delta = \frac{\ell(\pi^{(t)}) - \ell(\pi^{(t-1)})}{|\ell(\pi^{(t)})|}$
10: **return** $\tilde{\pi} = \tilde{\pi}^{(t)}$

And note that in Algorithm 1, Step 9 it should always be the case that $\ell(t) \geq \ell(t-1)$.

We note that this application of the EM algorithm is a very simple example of the standard EM, which in general does not guarantee a global maximum. However, this particular log-likelihood function (a sum of logarithms of weighted sums) is a concave function of its parameters, the $\pi_c$, and concavity ensures that any local maximum is a global maximum. In particular, we can ensure that there is a global, finite maximum and that the EM finds it by including a uniform component and making the algorithm start with all nonzero weights.
